# Amino acids trigger MDC-dependent mitochondrial remodeling by altering mitochondrial function

**DOI:** 10.1101/2024.07.09.602707

**Authors:** Nidhi Raghuram, Adam L. Hughes

## Abstract

Cells utilize numerous pathways to maintain mitochondrial homeostasis, including a recently identified mechanism that adjusts the content of the outer mitochondrial membrane (OMM) through formation of OMM-derived multilamellar domains called mitochondrial-derived compartments, or MDCs. MDCs are triggered by perturbations in mitochondrial lipid and protein content, as well as increases in intracellular amino acids. Here, we sought to understand how amino acids trigger MDCs. We show that amino acid-activation of MDCs is dependent on the functional state of mitochondria. While amino acid excess triggers MDC formation when cells are grown on fermentable carbon sources, stimulating mitochondrial biogenesis blocks MDC formation. Moreover, amino acid elevation depletes TCA cycle metabolites in yeast, and preventing consumption of TCA cycle intermediates for amino acid catabolism suppresses MDC formation. Finally, we show that directly impairing the TCA cycle is sufficient to trigger MDC formation in the absence of amino acid stress. These results demonstrate that amino acids stimulate MDC formation by perturbing mitochondrial metabolism.

**SUMMARY:** Raghuram and Hughes uncover a mechanism by which amino acids promote mitochondrial membrane remodeling via the Mitochondrial Derived Compartment (MDC) pathway. They show that elevated amino acids trigger MDC formation through a depletion of TCA cycle metabolites.

## INTRODUCTION

Mitochondria are double-membrane organelles that function as hubs for cellular metabolism and signaling (Chandel, 2014; Labbé et al., 2014). They play important roles in ATP production, heme and iron-sulfur cluster synthesis, amino acid, nucleotide and lipid metabolism, and regulation of metabolic, immune, and cell death signaling (Chandel, 2014; Epstein et al., 2001; Rutter and Hughes, 2015; Spinelli and Haigis, 2018). Mitochondria are highly dynamic, which allows their proper positioning at places of high energetic demand within cells and facilitates their adaptation to stress and changes in metabolic fuel sources (Alan and Scorrano, 2022; Labbé et al., 2014; Youle and Van Der Bliek, 2012). Dysregulated mitochondrial function has been linked to many age-related and metabolic disorders (Nunnari and Suomalainen, 2012; Wallace, 2005). Under conditions of stress or metabolic adaption, cells make use of multiple homeostasis mechanisms to optimize mitochondrial function. For example, mitophagy (Pickles et al., 2018), mitochondrial proteases (Quirós et al., 2015), the ubiquitin-proteasome system (Karbowski and Youle, 2011; Ravanelli et al., 2020), mitochondrial fission and fusion (Labbé et al., 2014), the mitochondrial unfolded protein response (Shpilka and Haynes, 2018), mitochondrial-derived vesicles (MDVs) (König and McBride, 2024; Sugiura et al., 2014), mitochondrial-derived compartments (MDCs) (Hughes et al., 2016; Schuler et al., 2021; Wilson et al., 2023a), and structures positive for outer membrane (SPOTs) (Li et al., 2022), all have roles in maintaining mitochondrial function and health. Failure of these mechanisms described above can be toxic to the cells and is implicated in a host of diseases (Nunnari and Suomalainen, 2012; Wallace, 2005).

Amongst these systems, we are focused on the recently discovered MDC pathway, which can remodel the mitochondria in response to changes in amino acids, mitochondrial phospholipids, or protein overabundance stress in the outer mitochondrial membrane (OMM) (Hughes et al., 2016; Schuler et al., 2021; Wilson et al., 2023a; Xiao et al., 2024). MDCs are dynamic multi-lamellar structures generated from OMM extensions that repeatedly elongate, invaginate and coalesce to ultimately form concentric spherical compartments that enclose cytosol (Wilson et al., 2023a, 2023b). They specifically sequester proteins localized in the OMM, whereas proteins in the matrix, inner mitochondrial membrane (IMM) as well as inter membrane space (IMS) are excluded (Hughes et al., 2016; Wilson et al., 2023a). MDCs form at sites of contact with the ER, and can be released from the organelle through fission and are degraded by autophagy (English et al., 2020; Hughes et al., 2016). While the exact function of MDCs remains undefined, current data suggest they provide mitochondria with the ability to selectively and dynamically adjust OMM content in response to a variety of conditions, including changes in amino acid and lipid abundance, as well as OMM protein load.

A major question regarding MDCs is how their formation is triggered by such apparently distinct stressors. While two types of MDC inducers (changes in OMM proteins and mitochondrial phospholipids) directly impact the organelle, it is unclear whether elevated intracellular amino acids also impact mitochondria directly, or whether they stimulate MDC formation through a mitochondrial-independent mechanism. In this study, we investigate how amino acids activate the MDC pathway. We provide evidence that MDCs are responsive to a mitochondrial signal that is triggered by amino acid elevation. Specifically, we find that excess amino acids create a strain on the TCA cycle, thus altering mitochondrial metabolic state to initiate MDC biogenesis. Based on this data, we propose that elevated amino acids, much like changes in mitochondrial lipids and protein content, trigger MDC formation by directly impacting the organelle.

## RESULTS

### Cellular growth state is important for MDC biogenesis

MDCs appear as foci forming from mitochondria that are enriched in OMM proteins such as Tom70, but exclude inner mitochondrial membrane (IMM) proteins such as Tim50 (Fig 1A) (Hughes et al., 2016). We previously showed that MDCs form in response to several stressors that cause a buildup in cytoplasmic amino acid pools in *S. cerevisiae*, including pharmacological- or age-induced dissipation of the proton gradient across the vacuole membrane, which prevents storage of amino acids in the vacuole, or impairment of protein translation, which precludes incorporation of amino acids into proteins and thus increases cytoplasmic pools (Hughes et al., 2016; Schuler et al., 2021). However, the mechanism by which amino acids activate the MDC pathway is unclear. While investigating the connection between amino acids and MDCs, we observed that in addition to amino acids, other growth cues influence the MDC pathway. Specifically, we found that MDC formation induced by the Vacuolar H^+^-ATPase (V-ATPase) inhibitor concanamycin A (concA) or the mTOR inhibitor rapamycin (rap) only occurs when yeast cells are cultured in a logarithmic phase of growth for an extended period of time. To induce MDCs, we culture cells overnight in glucose containing medium until they reach growth saturation the next day. At this point, we dilute the culture and grow the cells overnight again, keeping them at a low density to ensure log-phase growth. We found that while we observe MDCs in rap or concA-treated cells cultured for 16 hours in log-phase, MDC formation was blocked when cells were in saturation (0 hours of log-phase growth), and severely blunted when cells were cultured for only 3 hours in log-phase prior to concA or rap treatment (Fig 1A and B). The changes in MDC abundance strongly corelated with the physical appearance of the mitochondrial network. At 0-hours post saturation, the mitochondrial network was elaborate with many branches, indicative of mitochondria with enhanced respiratory activity (Westermann, 2012) (Fig 1A). At 3 hours in log-phase, mitochondria were still highly branched and elaborate, indicating that the mitochondria were still in a respiratory state at this point of growth (Fig 1A). After 16- hours of log-phase growth, the mitochondrial network was minimized with few branches, a phenotype that is typically observed in cells that are primarily utilizing fermentation (Fig 1A). These data suggest that MDC induction by concA and rap is impacted by the growth state of the cells.

**Figure 1.**
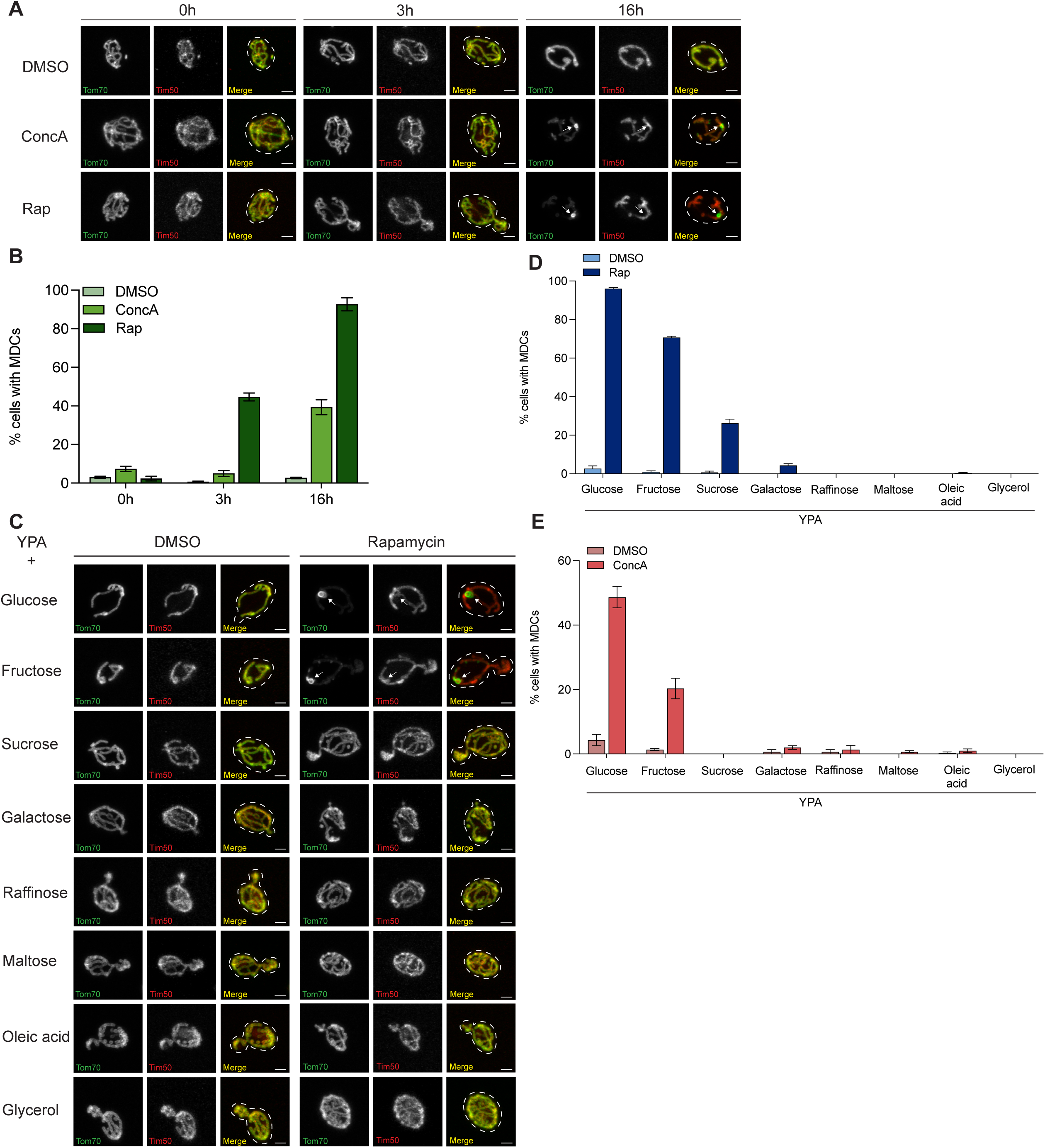
Cellular growth and respiratory state are important for MDC Biogenesis. (A) Super-resolution images of yeast cells grown to saturation overnight and diluted in fresh media for 0h, 3h or 16h, and then treated with concA or rap for 2h. Yeast cells are expressing Tom70-GFP and Tim50-mCherry. White arrow indicates MDC. Scale bar, 2μm. (B) Quantification of (A) showing the percentage of cells with MDCs. Error bars show Standard Error of Mean (+/- SEM) for N=3 replicates with n=100 cells per replicate. (C) Super-resolution images of yeast cells grown in high amino acid media (YPA) containing different carbon sources and treated with rap as indicated. Yeast cells are expressing Tom70-GFP and Tim50-mCherry. White arrow indicates MDC. Scale bar, 2μm. (D)Quantification of rap-induced MDC formation in cells grown in high amino acid (YPA) media containing varying carbon sources as indicated. Error bars show +/- SEM for N=3 replicates with n=100 cells per replicate. (E) Quantification of concA-induced MDC formation in cells grown in high amino acid (YPA) media containing varying carbon sources as indicated. Error bars show +/- SEM for N=3 replicates with n=100 cells per replicate.

### Carbon catabolite repression impacts MDC formation

Because we found that MDCs only formed when cells appeared to be in a fermentative state of growth, we wanted to directly assess the impact of respiration and fermentation on the MDC pathway*. S. cerevisiae* are Crabtree positive, meaning they prefer fermentation over respiration to produce energy. In the presence of fermentable sugars, such as glucose, yeast repress transcription of genes involved in mitochondrial respiration (De Deken, 1966). When fermentable sugars are absent from the growth medium, repression of respiratory genes is lifted, TCA cycle and electron transport chain (ETC) genes are transcribed, and the cells enter a respiratory metabolic state (De Deken, 1966; Klein et al., 1998). We tested whether the type of carbon source present in the growth medium impacted MDC formation. We performed MDC assays on yeast cells cultured for 16 hours post-saturation in media containing fermentable non-respiratory carbon sources—glucose or fructose. We observed robust MDC formation with concA and rap treatment for both sugars, although a small decrease in MDC formation was found in fructose grown cells. (Fig 1C-E). We observed tubular mitochondria with simple branching in cells grown in fermentable sugars as reported previously and similar to cells in mid log-phase growth (Fig 1C) (Egner et al., 2002). In contrast, in media containing sugars that are fermentable but do not repress mitochondrial respiration (sucrose, galactose, and raffinose), MDC formation was largely blunted. For rap treatment, 26% cells formed MDCs with sucrose supplementation while MDC formation was inhibited in galactose and raffinose-supplemented media (Fig 1D). A similar trend was observed for concA treated cells, where MDC formation was blocked in all three fermentable, respiratory sugars tested (Fig 1E). The mitochondria of these cells had increased branching, typical of cells in a respiratory state (Fig 1C). Finally, we observed a complete block in MDC formation with both concA and rap when cells were grown in media containing non-fermentable respiratory carbon sources—oleic acid, maltose, and glycerol (Fig 1C-E). Mitochondria in these cells were reticulate with elaborate branching, similar to what we observed in saturation (Fig 1A), and what is typical for mitochondria in a respiratory state. These results indicate that MDCs only form when cells are cultured in the presence of carbon sources that suppress mitochondrial biogenesis.

### Genetically stimulating mitochondrial biogenesis blocks MDC formation

Next, we wanted to test whether the carbon source-dependent effects on MDCs were caused by changes in mitochondrial respiratory capacity, or whether their impact was independent of mitochondrial metabolism. To do this, we overexpressed *HAP4* in yeast cells. Hap4 is a key transcriptional regulator of respiratory genes and is under the control of the glucose repression pathway (Klein et al., 1998). In glucose-containing medium, *HAP4* is repressed and thus inhibits expression of genes in the TCA cycle and oxidative phosphorylation. Therefore, by overexpressing *HAP4* in glucose-containing media, we can stimulate mitochondrial respiration when it would otherwise be repressed (Fig 2A) (Lascaris et al., 2002). To confirm the upregulation of the TCA cycle and ETC, we performed RNA sequencing analysis on cells overexpressing *HAP4* grown in glucose. Consistent with previous reports (Lascaris et al., 2002), we found that transcripts of genes involved in the TCA cycle and oxidative phosphorylation were upregulated in *HAP4* overexpressing cells compared to empty-vector (EV) control, despite being grown in glucose-containing media (Fig 2B-C). The mitochondria of *HAP4* overexpressing cells were reticulate with extensive branching, very similar to cells grown in non-fermentable carbon sources, and indicative of mitochondria with increased respiratory capacity (Fig 2D). We next tested the effect of overexpressing *HAP4* on MDC formation. MDC formation in an EV-control strain was normal, with 48% cells forming MDCs with concA treatment and 93% of cells forming MDCs with rap treatment. In contrast, *HAP4* overexpression blocked concA- and rap-induced MDC formation, even though the cells were grown in a fermentable carbon source (Fig 2E). This result indicates that stimulating mitochondrial activity blocks amino acid-induced MDC formation.

**Figure 2.**
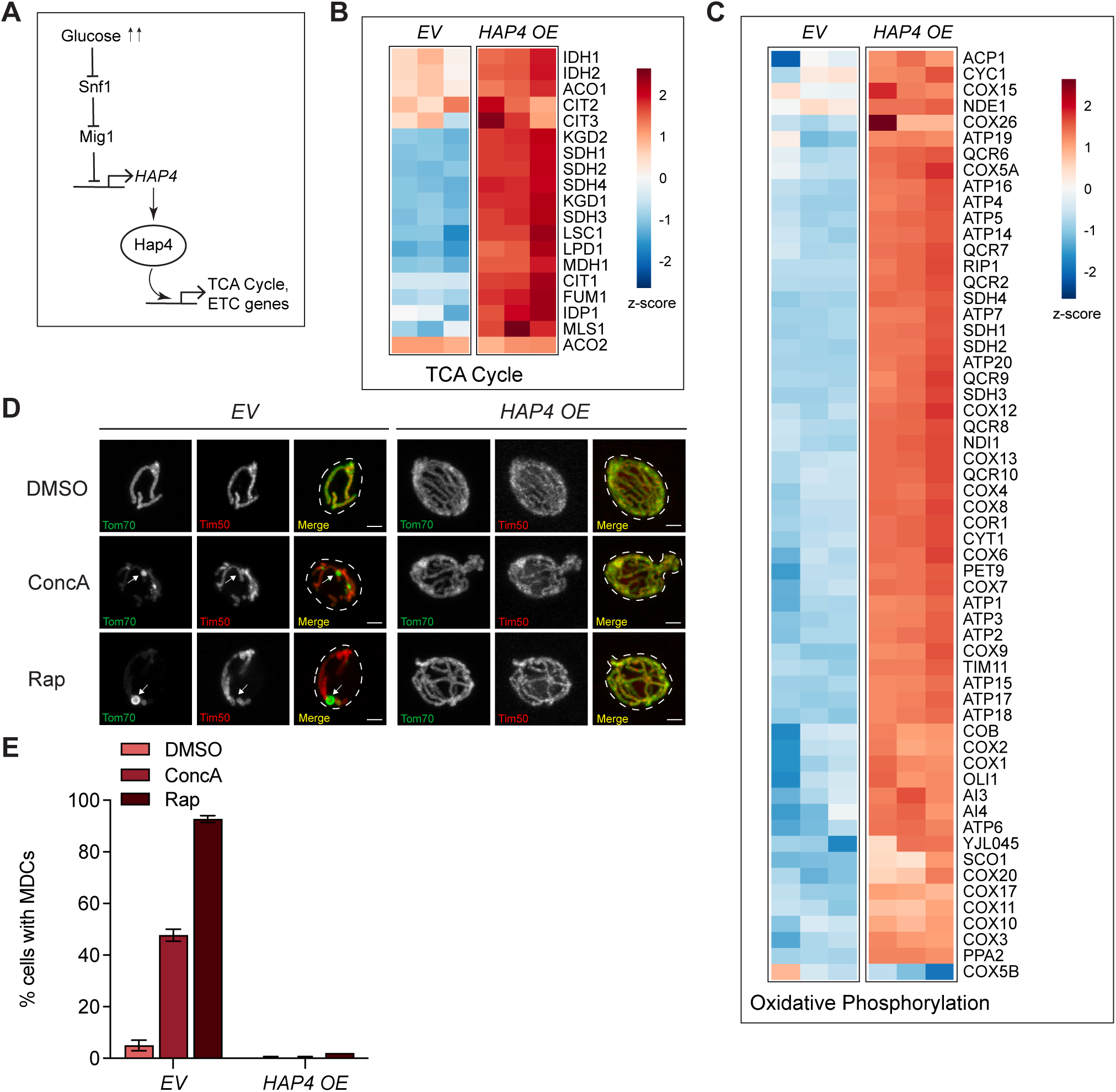
Genetically stimulating mitochondrial biogenesis blocks MDC formation. (A) Schematic showing glucose repression of *HAP4* and further regulation of TCA cycle and electron transport chain (ETC) genes by *HAP4*. (B) Heat maps showing z-scores of transcripts for TCA cycle genes from RNA sequencing analysis conducted on yeast strains overexpressing *HAP4*. Red=upregulated, Blue=downregulated. (C) Heat maps showing z-scores of transcripts for genes in the oxidative phosphorylation pathway from RNA sequencing analysis conducted on yeast strains overexpressing *HAP4*. Red=upregulated, Blue=downregulated. (D) Super-resolution images of *HAP4* overexpressing or *empty vector (EV)* control cells treated with concA or rap. Yeast cells are expressing Tom70-GFP and Tim50- mCherry. White arrow indicates MDC. Scale bar, 2μm. (E) Quantification of (D) showing the percentage of cells with MDCs. Error bars show +/- SEM for N=3 replicates with n=100 cells per replicate.

### Respiratory state does not impact plasma membrane nutrient transporter remodeling

We next wanted to determine how increasing mitochondrial biogenesis suppressed MDC formation. We first considered the possibility that amino acid elevation caused by concA or rap may be altered in respiratory carbon sources or growth states. To test this, we examined the influence of growth state, carbon source, and *HAP4* overexpression on the ability of rap and concA to activate another well-characterized amino acid sensitive system—removal of nutrient transporters from the plasma membrane through the ESCRT-dependent multivesicular body (MVB) pathway. It is well documented that elevated amino acids, triggered by impairment of protein translation and amino acid storage in vacuoles, stimulates ubiquitin-dependent removal of amino acid transporters from the PM and delivery to the vacuole for destruction (Hatakeyama and De Virgilio, 2019; Lin et al., 2008; Nikko and Pelham, 2009; Schuler et al., 2021). Moreover, our recent studies suggest a close coordination between the ESCRT/MVB pathway, vacuoles, and MDCs. ESCRT/MVB-dependent removal of PM transporters after concA or rap treatment occurs at the same time as MDC formation, and combined loss of vacuole amino acid storage, MDCs, and MVBs leads to amino acid-dependent cell death (Schuler et al., 2021). Interestingly, we found that in contrast to MDCs, removal of a representative GFP-tagged PM-localized amino acid transporter, Bap2-GFP (Grauslund et al., 1995), from the cell surface was not impacted by growth state, carbon source, or *HAP4* overexpression (Fig S1 A-C). Specifically, Bap2 was removed from the cell surface as assessed by line scan analysis upon treatment with concA, rap, and cycloheximide (CHX—another MDC inducer) in all conditions examined. This included at 0 and 3 hours post saturation (Fig S1A), after growth in the nonfermentable carbon source glycerol (Fig S1B), and in cells overexpressing *HAP4* (Fig S1C). Thus, in contrast to MDCs, amino acid-induced removal of PM transporters is unaffected by mitochondrial state. Importantly, this result indicates that an elevated amino acid signal is still present in rap and concA-treated cells grown in these different conditions and suggests that enhancing mitochondrial activity somehow prevents amino acid signaling to the MDC pathway.

### Elevated amino acids cause depletion of TCA cycle intermediates

To further understand how mitochondrial function and MDC formation could be linked, we considered the possibility that elevated amino acids may impact MDC formation by altering the TCA cycle. Unlike mammalian cells, many amino acids are removed from yeast cells through a catabolic route called the Ehrlich pathway, which involves conversion of amino acids into glutamate through transamination, followed by further metabolism to fusel aldehydes and alcohols which are released from the cell (Hazelwood et al., 2008). Using glucose tracing metabolic flux-analysis, we previously showed that during concA and rap treatment, the rate of glucose incorporated into glutamate via the TCA cycle metabolite-dependent transamination route shown in Fig. 3A was elevated (Schuler et al., 2021). We hypothesized that this may lead to a high demand on the TCA cycle intermediate α- ketoglutarate (α-kg) to support transamination reactions that function as the first step of amino acid catabolism via the Ehrlich pathway. Consistent with this hypothesis, we found that the steady-state abundance of TCA cycle intermediates immediately upstream and downstream of these transamination reactions, specifically α-ketoglutarate (α-kg) and succinic acid, respectively, declined after concA or rap treatment (Fig 3 B-C). Cells grown in the non-fermentable carbon source glycerol had higher levels of the TCA cycle intermediates α-ketoglutarate and succinic acid compared to cells grown in glucose media. Moreover, the levels of these metabolites remained higher than untreated glucose grown cells upon treatment with concA and rap (Fig 3B-C). A similar trend was observed in cells with *HAP4* overexpression, where levels of α-kg and succinic acid were higher compared to the EV control and their levels remained largely unaffected after concA treatment (Fig 3D).

**Figure 3.**
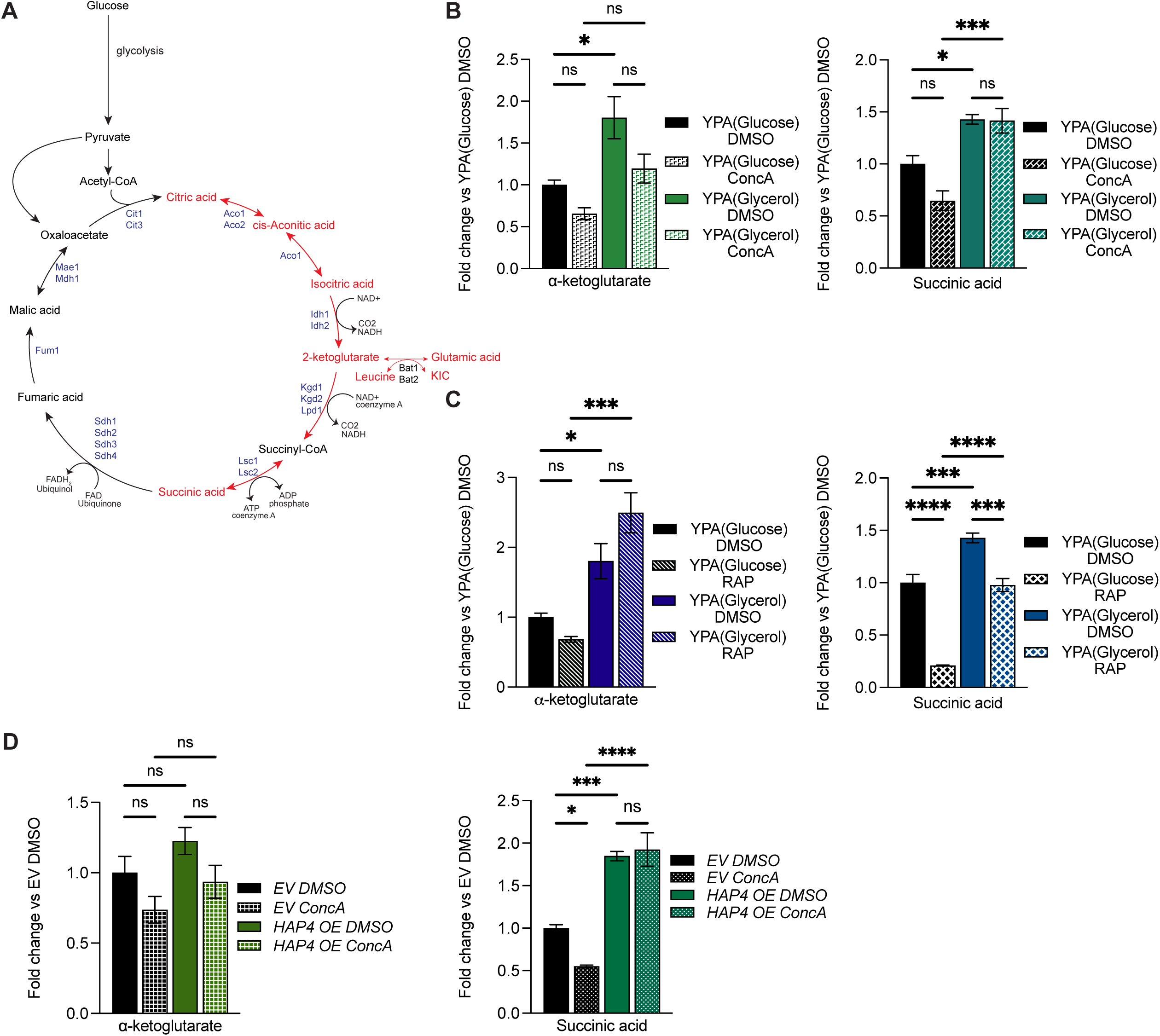
Elevated amino acids cause depletion of TCA cycle intermediates in yeast. (A) Schematic of cellular glucose, TCA cycle, amino acids metabolism and transamination reaction. (B) Analysis of whole cell α-kg and succinic acid metabolite levels in yeast cells grown in high amino acid media (YPA) containing glucose or glycerol and treated with concA for 2 hours as indicated. Error bars show +/- SEM for N=4 replicates. Statistical comparison shows difference to the corresponding DMSO control and difference to the corresponding glucose control. n.s., not significant, *p<0.0332, ***p<0.0002, One- way ANOVA with Šídák’s multiple comparisons test. (C) Analysis of whole cell α-kg and succinic acid metabolite levels in yeast cells grown in high amino acid media (YPA) containing glucose or glycerol and treated with rap for 2 hours as indicated. Error bars show +/- SEM for N=4 replicates. Statistical comparison shows difference to the corresponding DMSO control and difference to the corresponding glucose control. n.s., not significant, *p<0.0332, ***p<0.0002, ****<0.0001, One-way ANOVA with Šídák’s multiple comparisons test. (D) Analysis of whole cell α-kg and succinic acid metabolite levels in *EV* and *HAP4 OE* cells treated with concA for 2 hours. Error bars show +/- SEM for N=4 replicates. Statistical comparison shows difference to the corresponding DMSO control and difference to the corresponding *EV* control. n.s., not significant, *p<0.0332, ***p<0.0002, ****<0.0001, One-way ANOVA with Šídák’s multiple comparisons test.

Altogether, these results suggest that elevated levels of amino acids caused by treatment with concA and rap leads to a reduction in two TCA cycle intermediates, and that this depletion is blocked in non-fermentable carbon sources or in cells with genetically enhanced mitochondrial biogenesis.

### Increasing TCA cycle activity inhibits MDC formation

Based on our results, we hypothesized that a disruption in the TCA cycle or an elevated utilization of TCA cycle metabolites to support amino acid catabolism may be what ultimately triggers MDC formation in the presence of high amino acids. To test this hypothesis, we used two independent approaches to boost TCA cycle metabolite levels and assessed the impact on MDC formation. First, we genetically manipulated the so-called retrograde (RTG) pathway, which controls transcription of several genes in the TCA cycle and glyoxylate cycle during growth in fermentable carbon sources. An important function of the retrograde pathway is to maintain sufficient levels of α-kg by increasing expression of TCA cycle and glyoxylate pathway enzymes in order to support glutamate production during times of mitochondrial impairment. This pathway relies on the transcriptional activators Rtg1 and Rtg3, whose access to the nucleus is controlled by mitochondrial functional state via a signaling cascade that involves Mks1, a negative regulator of the pathway (Fig 4A) (Butow and Avadhani, 2004; Jazwinski and Kriete, 2012; Liu and Butow, 2006). Deletion of the negative regulator Mks1 leads to nuclear localization of Rtg1 and Rg3, causing constitutive activation of the pathway and elevated expression of retrograde target genes in the presence of fermentable carbon sources. In contrast, deletion of Rtg1 or Rtg3 impairs the pathway and lowers expression of TCA and glyoxylate cycle enzymes. We performed RNA sequencing analysis on cells lacking *MKS1* or *RTG1*, and confirmed previous observations that TCA cycle and glyoxylate cycle genes were highly upregulated in *mks1*1Δ** cells and significantly downregulated in *rtg1*1Δ** cells compared to wildtype (Fig 4B) (Epstein et al., 2001). Furthermore, steady state metabolite analysis on *mks1*1Δ** cells showed that the levels of TCA cycle metabolites, α-kg and succinic acid, were significantly higher as compared to the wildtype control, and remained elevated upon treatment with concA (Fig 4C). Importantly, we assessed MDC formation in *mks1*1Δ** cells and found that, compared to wildtype, MDC activation was blunted. Only 25% of *mks1*1Δ** cells formed MDCs upon concA treatment, as compared to 50% observed in wild-type cells. Similarly, while 90% of wildtype cells formed MDCs with rap treatment only 53% of *mks1*1Δ** mutant cells formed MDCs (Fig 4D-E). These results suggest that increasing TCA and glyoxylate cycle enzymes and metabolite levels can partially block MDC formation.

**Figure 4.**
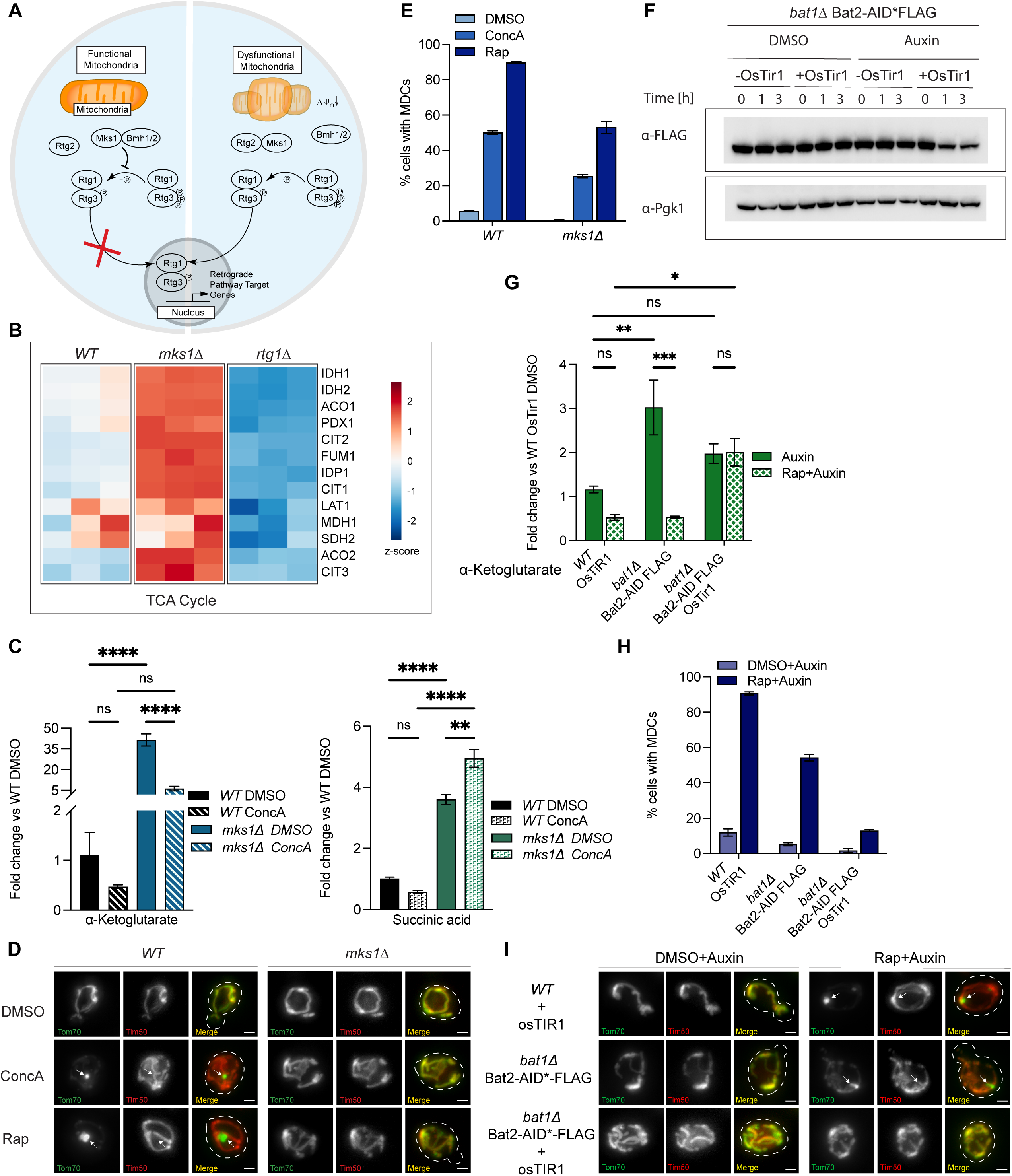
Increasing TCA cycle activity inhibits MDC formation. (A) Schematic showing activation of the retrograde pathway and its target genes under conditions of mitochondrial dysfunction, linking the mitochondria and nucleus. (B) Heat map showing z-score of transcripts from RNA sequencing analysis conducted on *WT, mks1Δ* and *rtg1Δ* yeast strains. Red=upregulated, Blue=downregulated. Gene function indicated. (C) Analysis of whole cell α-kg and succinic acid metabolite levels in *WT* and *mks1Δ* mutant cells treated with concA for 2 hours. Error bars show +/- SEM for N=3 replicates. Statistical comparison shows difference to the corresponding DMSO control and difference to the corresponding *WT* control. n.s., not significant, **p<0.0021, ****<0.0001, One-way ANOVA with Šídák’s multiple comparisons test. (D) Widefield images of *WT* and *mks1Δ* mutant cells treated with concA or rap. Cells are expressing Tom70-GFP and Tim50-mCherry. White arrow indicates MDC. Images show maximum intensity projections. Scale bar, 2μm. (E) Quantification of (D) showing the percentage of cells with MDCs. Error bars show +/- SEM for N=3 replicates with n=100 cells per replicate. (F) Western blot showing time course of auxin induced Bat2-AID*-FLAG depletion in the presence or absence of OsTir1. (G) Analysis of whole cell α-kg metabolite levels in *WT*+OsTir1 and indicated mutant cells pretreated with auxin for 1h followed by DMSO+auxin or rap+auxin treatment for 2 hours. Error bars show +/- SEM for N=3 replicates. Statistical comparison shows difference to the corresponding DMSO control and difference to the corresponding *WT* control. n.s., not significant, *p<0.0332, **p<0.0021, ***p<0.0002, Two-way ANOVA with Tukey test. (H) Quantification of MDC formation in *WT*+OsTir1 and indicated mutant cells pretreated with auxin for 1h followed by DMSO+auxin or rap+auxin treatment for 2 hours. Error bars show +/- SEM for N=3 replicates with n=100 cells per replicate. (I) Widefield images of *WT*+OsTir1 and indicated mutant cells pretreated with auxin for 1h followed by DMSO+auxin or rap+auxin treatment for 2 hours. Cells are expressing Tom70-GFP and Tim50-mCherry. White arrow indicates MDC. Images show maximum intensity projections. Scale bar, 2μm.

In addition to manipulating the retrograde pathway, we also tested whether preventing usage of TCA metabolites to support amino acid catabolism by genetically disrupting transaminases would prevent MDC activation. Because elevated branched chain amino acids are a potent stimulator of MDC formation (Schuler et al., 2021), we conditionally eliminated branched chain amino acid transaminase activity and assessed the impact on TCA metabolites and MDC formation during amino acid stress. We did this by conditionally reducing the levels of the branched-chain transaminase Bat2 in cells deleted for the other branched chain transaminase Bat1. This was achieved by fusing Bat2 to an auxin inducible degron (AID) which targets the tagged protein for proteasomal degradation upon addition of auxin (Morawska and Ulrich, 2013). A reduction in FLAG-tagged Bat2 protein was achieved within 1h of auxin addition (Fig 4F). From steady state metabolite analysis, we observed higher levels of α-kg in cells lacking Bat1 alone (*bat1*1Δ** Bat2-AID FLAG without auxin-inducible E3 ligase OsTir1), as well as in *bat1*1Δ** cells with transient auxin-induced loss of Bat2 (*bat1*1Δ** Bat2-AID FLAG with OsTir1) (Fig 4G). α-kg levels were largely unaffected by rap treatment in *bat1*1Δ** Bat2-AID mutant cells in the presence of auxin, which is in contrast to wildtype and cells lacking Bat1 alone, where α-kg levels were decreased with rap (Fig 4G). Importantly, we found that rap treatment induced MDC formation in 91% of wildtype cells but only 54% of *bat1*1Δ** mutant cells, and 13% of *bat1*1Δ** Bat2-AID cells (Fig 4H-I). Collectively, these results indicate that maintaining levels of TCA cycle metabolites under conditions of elevated amino acid stress inhibits MDC biogenesis.

### Impairing the TCA cycle is sufficient to activate the MDC pathway

Finally, given that boosting the levels of the TCA cycle enzymes and metabolites can block MDC formation in the presence of high amino acids, we sought to determine whether impairment of the TCA cycle was sufficient to trigger MDC formation in the absence of amino acid stress. To do this, we again made use of genetic manipulation of the retrograde pathway. Specifically, we deleted *RTG1*, which lowers expression of TCA and glyoxylate cycle enzymes and reduces steady state levels of most TCA cycle metabolites (Fig 4B and Fig 5A). We found that deletion of *RTG1* caused constitutive MDC formation in 62% of cells, without any additional treatments or stressors (Fig 5B-C). Similarly, *rtg2*1Δ** and *rtg3*1Δ** mutants, which also have an impaired retrograde pathway, also exhibited constitutive MDC formation (Fig S2A). To confirm that the constitutive MDC formation observed was due to the deletion of *RTG1* and not due to any secondary effects, we overexpressed *RTG1* in *rtg1*1Δ** cells to rescue retrograde pathway function. We found that overexpressing *RTG1* restored MDC formation in *rtg1*1Δ** mutant similar to wildtype levels, where less than 5% of *rtg1*1Δ** mutant and wildtype cells formed MDCs in untreated cells (Fig S2B). Additionally, because deletion of *rtg1*1Δ** caused a depletion in the first half of the TCA cycle specifically (Fig 5A), we tested whether re-expressing enzymes in this part of the TCA pathway could retore MDC levels back to wildtype in the absence of *RTG1*. While we were unable to restore expression of every enzyme in this pathway, we found that combined overexpression of *CIT1* and *ACO1* (see Fig 3A) partially restored levels of TCA cycle metabolites (Fig S2C), and reduced MDC levels in *rtg1*1Δ** mutants back towards wild-type levels (Fig 5D).

**Figure 5.**
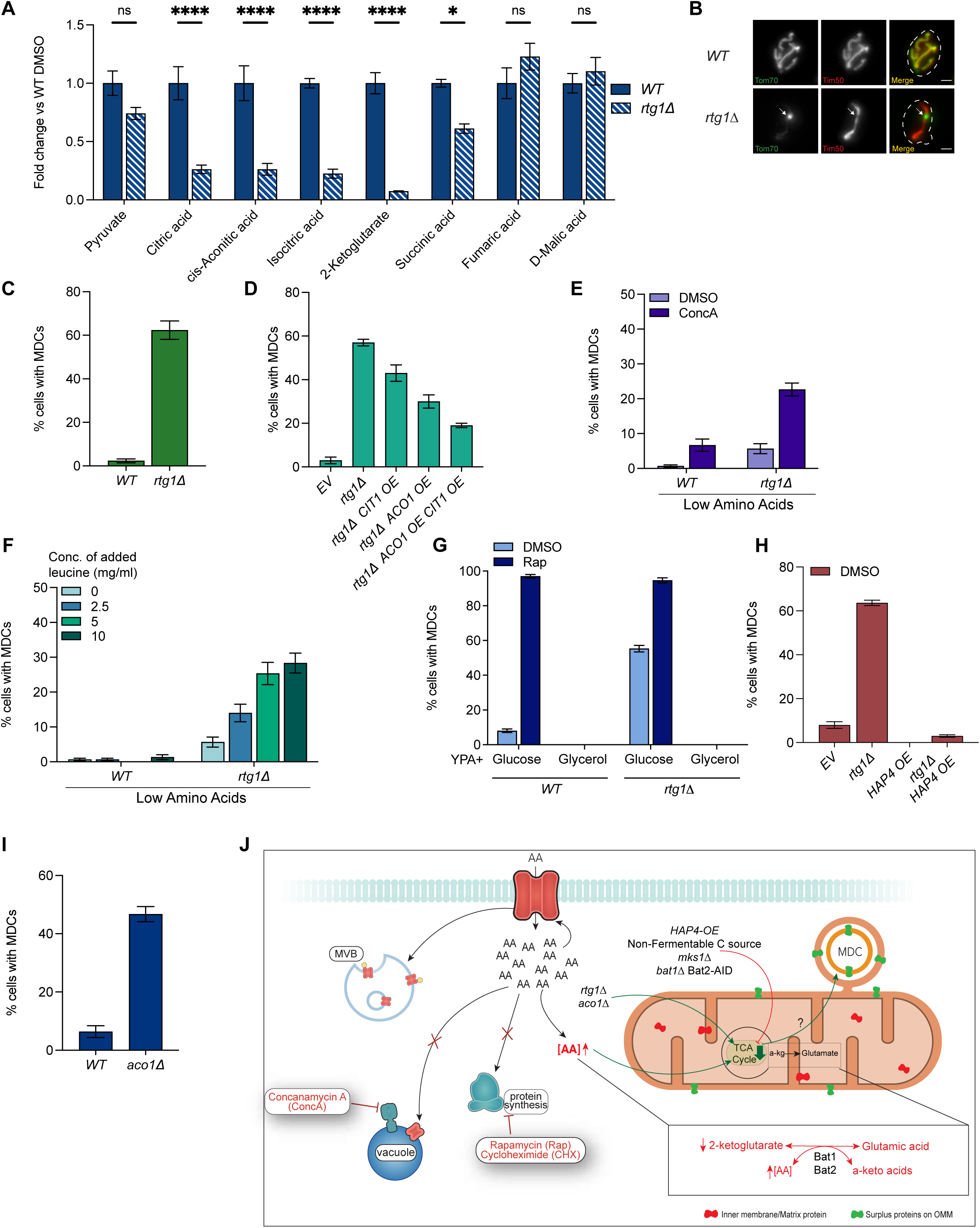
Impairing the TCA cycle is sufficient to activates the MDC pathway. (A) Analysis of whole cell pyruvate and TCA cycle metabolite levels in *WT* and *rtg1Δ* mutant cells. Error bars show +/- SEM for N=4 replicates. Statistical comparison shows difference to the corresponding *WT* control. n.s., not significant, *p<0.0332, ****<0.0001, Two-way ANOVA with Šídák’s multiple comparisons test. (B) Widefield images of *WT* and *rtg1Δ* mutant cells. Cells are expressing Tom70-GFP and Tim50-mCherry. White arrow indicates MDC. Images show maximum intensity projections. Scale bar, 2μm. (C) Quantification of MDC formation in *WT* and *rtg1Δ* mutant cells. Error bars show +/- SEM for N=3 replicates with n=100 cells per replicate. (D) Quantification of MDC formation in *EV* and indicated mutant cells. Error bars show +/- SEM for N=3 replicates with n=100 cells per replicate. (E) Quantification of concA-induced MDC formation in *WT* and *rtg1Δ* mutant cells, grown in low amino acid media. Error bars show +/- SEM for N=3 replicates with n=100 cells per replicate. (F) Quantification of MDC formation in *WT* and *rtg1Δ* mutant cells, grown in low amino acid media supplemented with 2.5, 5 or 10 mg/ml leucine. Error bars show +/- SEM for N=3 replicates with n=100 cells per replicate. (G) Quantification of MDC formation in *WT* and *rtg1Δ* mutant cells, grown in high amino acid media (YPA), containing glucose or glycerol and treated with rap as indicated. Error bars show +/- SEM for N=3 replicates with n=100 cells per replicate. (H) Quantification of MDC formation in *EV* and indicated mutant cells. Error bars show +/- SEM for N=3 replicates with n=100 cells per replicate. (I) Quantification of MDC formation in *WT* and *aco1Δ* mutant cells. Error bars show +/- SEM for N=3 replicates with n=100 cells per replicate. (J) Described left to right. Excess intracellular amino acids(AA) caused by concA, rap or cHX trigger internalization of PM transporters through the MVB pathway. Excess AA also trigger MDC pathway activation. Excess AA create a strain on the TCA cycle by utilizing the TCA cycle metabolite α-kg to support AA catabolism and activate the MDC pathway. Directly impairing the TCA cycle by deletion of *RTG1* or *ACO1* activates the MDC pathway. Preventing depletion of TCA cycle by deletion of *MKS1,* overexpression of *HAP4,* growth in nonfermentable carbon source or *bat1Δ* Bat2- AID*-FLAG OsTir1 inhibits MDC formation even in conditions of AA stress.

Because the presence of branched chain amino acids in yeast cells increases the demand on TCA metabolites to support amino acid catabolism, we wondered whether we could suppress MDC formation in *rtg1*1Δ** cells by lowering the amino acid catabolic burden. Indeed, we found that growth of *rtg1*1Δ** cells in low amino acid medium reduced MDC levels. MDCs formed in 6% *rtg1*1Δ** mutant cells in low amino acids medium, similar to levels in wild-type cells (Fig 5E). Moreover, we found that MDC levels were partially restored by elevating amino acids with concA in *rtg1*1Δ** mutants (Fig 5E) and that addition of leucine to low amino acid medium strongly restored MDC formation in *rtg1*1Δ** mutants (Fig 5F). These results suggest that in presence of amino acids, the retrograde pathway plays a critical role in maintaining TCA metabolite levels to support amino acid catabolism, and that removing the burden of amino acid catabolism suppresses MDC formation in the absence of a functional retrograde pathway. Interestingly, we also found that MDC formation in *rtg1*1Δ** mutants was suppressed by growing cells in non-fermentable carbon sources or overexpression of Hap4 (Fig 5G-H). These results are consistent with previous studies showing that the RTG pathway is a critical regulator of TCA cycle activity in glucose grown cells, but that Hap4 is the dominant regulator of these same genes when it is activated in non-fermentable carbon sources (Liu and Butow, 1999). Finally, we found that impairment of the TCA cycle by knocking out the conserved TCA cycle enzyme aconitase (Aco1) was also sufficient to stimulate MDC formation, similar to what we observed in cells lacking the retrograde pathway (Fig 5I). Collectively, these results suggest that impairment of the TCA cycle is sufficient to stimulate MDC formation in the absence of amino acid stress.

## DISCUSSION

Prior to this work, it was known that MDCs are cargo selective multilamellar domains that facilitate the removal of content from the OMM (Hughes et al., 2016; Wilson et al., 2023a, 2023b). These structures, which form at ER-mitochondrial contacts in cells, are triggered by several different perturbations in cells, including changes in mitochondrial protein and lipid content, as well as alterations in cellular amino acid levels (English et al., 2020; Schuler et al., 2021; Wilson et al., 2023a; Xiao et al., 2024). While the full function of MDCs remains unclear, a current working model is that they serve as a mechanism to specifically adjust the size and/or composition of the OMM in response to a variety of different cellular stresses.

Why this occurs in cells is unclear, but one speculative idea is that this may provide the organelle with a mechanism to help balance its two distinct membranes. Of the three stresses that are known to trigger MDCs, two of them (lipids and protein content changes) appear to directly alter some aspect of the mitochondria (Wilson et al., 2023a; Xiao et al., 2024). In contrast, how and why MDCs are triggered by elevated amino acids is unclear. In our current study, we sought to understand how amino acids regulate MDCs, and our results provide new insight into how these metabolites regulate the MDC pathway. Specifically, as depicted in Fig 5J, our data supports a model that excess amino acids in yeast, which occurs when pathways that control amino acid uptake, usage and localization are compromised, stimulates MDC biogenesis by causing a high demand on the TCA cycle to support transamination-based amino acid catabolism. Consistent with this idea, boosting TCA cycle metabolites and enzyme levels blocks amino-acid induced MDC formation, as does broad stimulation of mitochondrial biogenesis. Moreover, directly impairing the TCA cycle in the absence of amino acid stress is sufficient to stimulate MDC formation. Thus, it appears that amino acids, much like changes in lipids and protein load in the OMM, trigger MDC activation by altering a functional aspect of the organelle itself. Collectively, these results suggest that MDCs remodel mitochondria in response to changes in their structure and/or function.

Interestingly, our findings that MDCs respond to TCA cycle-related cues differentiates MDC signaling and activation from known mechanisms by which elevated amino acids post-translationally regulate the abundance of plasma membrane nutrient transporters through the ESCRT/MVB pathway (Lin et al., 2008; Nikko and Pelham, 2009). Thus, while elevated amino acids in cells can simultaneously trigger MDC formation and ESCRT/MVB-dependent nutrient transporter destruction, MDCs respond to the effects of amino acids on a functional aspect of the organelle. Importantly, unlike PM transporter removal, MDC formation can be mitigated when TCA cycle metabolite levels are maintained. These results highlight a previously unappreciated link between TCA cycle activity and outer OMM remodelling.

Interestingly, a connection between TCA cycle and changes in mitochondrial morphology has been identified before, wherein accumulation of the TCA cycle metabolite fumarate can lead to alterations in mitochondrial morphology through release of mitochondrial derived vesicles (MDVs). These MDVs are cargo selective for matrix proteins and contained mitochondrial DNA (Zecchini et al., 2023). Although we do not understand how impairment of the TCA cycle triggers MDC formation in the current study, it appears that cells may initiate various remodelling mechanisms to alter mitochondria in response to changes in TCA cycle activity. Thus, our study adds to a growing list of examples of how cells alter mitochondrial structure in response to metabolic cues.

Finally, the discovery of an important role for mitochondrial metabolic state and TCA cycle activity in MDC formation raises several additional questions, including how changes in TCA cycle activity trigger MDC formation, and whether this mechanism is similar to MDC formation induced by changes in phospholipids and protein overload stress. At this point, it is unclear whether a particular TCA cycle metabolite is involved, or if changes in the TCA cycle alter another aspect of the organelle that stimulates MDC formation. In a previous study, we showed that amino acid-induced MDCs were triggered by a downstream decrease in mitochondrial PE levels (Xiao et al., 2024), and that PE decline alone was sufficient to induce MDCs. Our current data indicating an additional involvement of TCA cycle perturbations downstream of amino acid stress strongly suggests that an unknown link between the TCA cycle and mitochondrial phospholipid levels regulates the MDC pathway.

What this connection is and how it is regulated is currently unclear, but will be an important area of future investigation. In summary, while many unanswered questions remain, the results from this study provide an important step forward in our fundamental understanding of the MDC pathway and its role in mitochondrial homeostasis.

## MATERIALS AND METHODS

### Yeast strains

All yeast strains are derivatives of S288C (BY) (Baker Brachmann et al., 1998) and are listed in Table S1. Strains expressing fluorescently tagged *TOM70*, *TIM50, BAP2* and/or AID*- 6xFLAG tagged *BAT2* from their native loci were created using one-step PCR mediated C-terminal endogenous epitope tagging using standard techniques and oligo pairs listed in Table S2. Plasmid templates for fluorescent epitope tagging were from the pKT series of vectors (Sheff and Thorn, 2004). Plasmids used for AID*-6xFLAG tagging and integration of GPD-OsTir1 into the *URA3* locus are described below. Correct integrations were confirmed by a combination of colony PCR across the chromosomal integration site and correctly localized expression of fluorophore by microscopy. For strains bearing deletions of *RTG1* and *MKS1*, the gene was deleted in haploids of both mating types by one step PCR-mediated gene replacement with the indicated selection cassette using standard techniques and oligo pairs listed in Table S2. For strains bearing deletion for *ACO1*, one copy of the gene was deleted using the same method described above to generate a heterozygous diploid which was subsequently sporulated to get haploid mutants. The resulting haploid mutants were then mated to generate diploids. Plasmid templates for gene deletions were from the pRS series of vectors (Sikorski and Hieter, 1989). Correct insertion of the selection cassette into the target gene was confirmed by colony PCR across the chromosome insertion site. Yeast strains constitutively expressing *HAP4, CIT1* and *ACO1* under the control of a GPD promoter were generated by integration of expression cassette into yeast chromosome I (199456-199457). Plasmids for expression of GPD driven expression cassettes are described in Table S3. Correct integration of expression cassette into chromosome I was confirmed by colony PCR across the chromosomal insertion site.

Wild-type yeast strains AHY4706 and AHY11684 *HAP4* OE strain AHY11682, *rtg1*1Δ** strains AHY11253 and *mks1*1Δ** strain AHY11209 which were rendered prototrophic with pHLUM (see Table S3) to prevent complications generated by amino acid auxotrophies in the BY strain background, were used for gene expression analysis. Yeast strains AHY1480, AHY11574, AHY11572, AHY13782, AHY11037, AHY15993, AHY15482, AHY11132, AHY13023, AHY14599, AHY11933, AHY12201, AHY12753, AHY12755, AHY12757, AHY11446, AHY11448, AHY11626, AHY12944, AHY12946, AHY11624, AHY11626, AHY11160 were used for quantification of MDC formation using widefield microscopy. AHY1480, AHY11574 and AHY11572 were used for obtaining microscopy images using super-resolution microscopy. Metabolomics analysis was performed in AHY1480, AHY11572, AHY11574, AHY11626, AHY13782, AHY11933, AHY12201, AHY12753, AHY12755, AHY12757, AHY15593, AHY11037 and AHY15482. A complete list of strains used in this manuscript can be found in Table S1.

### Yeast cell culture and media

Yeast cells were grown exponentially for a period of 15-20 hours at 30°C to a maximum density of 3-8 x 10^6^ cells/ml before the start of all experiments described in the paper including MDC assays as well as sample preparation for RNA sequencing and metabolomics (except where indicated). Unless otherwise indicated, cells were cultured in YPAD (1% yeast extract, 2% peptone, 0.005% adenine, and 2% glucose) medium. For carbon source dependent experiments, cells were cultured in YPA (1% yeast extract, 2% peptone, 0.005% adenine) medium with the indicated carbon source added. Carbon sources were added to a final concentration of 2% except otherwise indicated. Oleic acid was added to a final concentration of 1mM with 1% tween. Glycerol was added to a final concentration of 3%. For experiments in low amino acid media, cells were cultured in SD (containing 0.67% yeast nitrogen base without amino acids, 2% glucose, supplemented nutrients 0.074 g/L each of adenine, alanine arginine, asparagine aspartic acid, cysteine, glutamic acid, glutamine, glycine, histidine, myo-inositol, isoleucine, lysine, methionine, phenylalanine, proline, serine, threonine, tryptophan, tyrosine, uracil, valine, 0.369 g/L of leucine and 0.007 g/L of para-aminobenzoic acid.) Concentrations used for drug treatments were compounds were: concanamycin A (500nM), rapamycin (200nM) and auxin (1mM) unless otherwise indicated.

### Plasmids

Plasmids used in this study are listed in Table S3. pHLUM, a yeast plasmid expressing multiple auxotrophic marker genes from their endogenous promoters, was obtained from Addgene (#40276) (Mülleder et al., 2012). pHYG-AID*-6FLAG (Morawska and Ulrich, 2013) and pGPD1-osTIR1-URA3 (Parnell et al., 2023) were described previously. To integrate pGPD1-osTIR1 into the *URA3* locus, pGPD1-osTIR1-URA3 was digested by Pme1. Plasmids for GPD driven expression of *HAP4, CIT1, ACO1, RTG1* were generated by gateway mediated transfer of corresponding ORF (Harvard Institute of Proteomics) from pDONR 201/221 into a pAG306-ccdB chromosome I (Hughes and Gottschling, 2012) using Gateway LR Clonase II enzyme mix (ThermoFisher) according to the manufacturer’s instructions. To integrate the resulting expression plasmid into yeast chromosome I (199456- 199457), pAG306GPD-ORF chromosome I was digested with NotI. All insert sequences were verified by Azenta/Genewiz sequencing.

### Yeast MDC Assays

MDC assays were performed as described previously (Schuler et al., 2021). In brief, unless otherwise indicated, overnight log phase grown cell cultures were treated with vehicle, dimethyl sulfoxide (DMSO) or indicated drugs for two hours. For assays testing the effect of time in log phase on MDC formation, cells were diluted into fresh media for the indicated amount of time, following which, the cells were grown in the presence of DMSO or indicated drugs for two hours. After two hours incubation with vehicle or drugs, cells were pelleted by centrifugation, resuspended in imaging buffer (5% glucose, 10mM HEPES pH 7.6) and imaged as described below.

### Microscopy

200nm optical z sections of live yeast cells were acquired with an AxioObserver (Carl Zeiss) equipped with a Zeiss Axiocam 506 monochromatic camera and 63 x oil immersion objectives (Carl Zeiss, Plan Apochromat, NA 1.4) for quantification of MDC formation. Single z planes for images showing plasma membrane (PM) transporter internalization were acquired with AxioImager M2 (Carl Zeiss) equipped with an edge 4.2 CMOS camera (PCO) and 63 x oil immersion objective (Carl Zeiss, Plan Apochromat, NA 1.4). Widefield images were acquired with Zen (Carl Zeiss) and processed in Fiji (Schindelin et al., 2012). Super-resolution images showing varied mitochondrial morphology and MDC formation in live yeast cells were acquired with an LSM800 (Carl Zeiss) equipped with an Airyscan detector and 63 x oil-immersion objective (Carl Zeiss, Plan Apochromat, NA 1.4).

### Image Analysis

Super-resolution images were acquired with Zen (Carl Zeiss), processed using the Airyscan processing algorithm on Zen (Carl Zeiss) and Fiji. Individual channels of all images were minimally adjusted in Fiji to match the fluorescence intensity between channels for better visualization. Unless specified, line-scan analysis for PM transporter internalization was performed on non-adjusted single z sections from widefield images. Line scan analysis for PM transporter internalization assays on cells grown to varied time in log phase, was performed on minimally adjusted z sections of widefield images to remove background.

Maximum intensity projected widefield images generated in Zen (Carl Zeiss) were used to quantify the percentage of cells with MDCs. MDCs were identified as structures of varying size and shape, enriched in Tom70 and negative in Tim50. Unless specified, maximum intensity projected images are displayed for all yeast images.

### RNA Isolation and Sequencing

4 x 10^7^ of overnight log-phase yeast cell cultures were harvested and the pellets were snap frozen and stored at -80 °C. The cells were then lysed via micro-bead beating in Omni Bead Ruptor 12 Homogenizer, and total RNA was enriched using RNeasy mini kit (Qiagen).

Contaminating DNA was removed by on column RNase-free DNase (Qiagen) treatment. Following RNA isolation, RNA sequencing was performed with the High-Throughput Genomics and Bioinformatics Analysis Shared Resource at Huntsman Cancer Institute at the University of Utah. Total RNA samples (100ng-500ng) were hybridized with Ribo-Zero Gold to substantially deplete rRNA from the samples. Stranded RNA sequencing libraries were prepared using Illumina TruSeq Stranded Total RNA Library Prep Gold kit (20020598) with TruSeq RNA UD Indexes (20022371). Purified libraries were qualified on an Agilent Technologies 4150 Tape Station using a D1000 ScreenTape assay. The molarity of adapter- modified molecules was defined by quantitative PCR using a Kapa Biosystem Kapa Library Quant Kit. Individual libraries were normalized to 1.30 nM in preparation for illumina sequence analysis. Sequencing libraries were chemically denatured and applied to an Illumina NovaSeq flow cell using the NovaSeq XP workflow (20043131). Following transfer of the flowcell to an Illumina NovaSeq 6000 instrument, a 150 x 150 cycle paired end sequence run was performed using a NovaSeq 6000 S4 reagent Kit v1.5 (20028312).

### RNA Sequencing Bioinformatics

The yeast R64-1-1 genome and gene annotation files were downloaded from Ensembl release 102. Missing UTRs in the gene annotation file were added using microarray data from (Xu et al., 2009) and saved to SacCer3_R64_all_genes_NoDubious_UTR.nochr.gtf. Optical duplicates were removed from the paired end FASTQ files using clumpify v38.34 and reads were trimmed of adapters using cutadapt v2.8 (Martin, 2011). The trimmed reads were aligned to the yeast reference database using STAR version 2.7.6a in two pass mode to output a BAM file sorted by coordinates (Dobin et al., 2013). Mapped reads were assigned to annotated genes using featureCounts version 1.6.3 (Liao et al., 2014). The output files from cutadapt, FastQC, FastQ Screen, Picard CollectRnaSeqMetrics, STAR and featureCounts were summarized using MultiQC to check for any sample outliers (Ewels et al., 2016). Differentially expressed genes were identified using a 5% false discovery rate with DESeq2 version 1.30.1 (Love et al., 2014). Significant genes were analysed using the YeastEnrichr website (Chen et al., 2013) to find over-represented pathways and GO terms. All RNA sequencing data is provided in Table S4.

### Protein Preparation and Immunoblotting

For western blot analysis of protein levels, overnight log phase grown cells were treated as indicated, 2x10^^7^ cells were harvested, and cell pellets were shock frozen in liquid nitrogen. Subsequently, the cell pellets were washed with dH2O and incubated in 0.1M NaOH for five minutes at RT. The cells were reisolated by centrifugation at 14000g for ten minutes at 4°C following which, the cells were lysed for five minutes at 95°C in lysis buffer (10 mM Tris pH 6.8, 100 mM NaCl, 1 mM EDTA, 1 mM EDTA, 1% SDS). Samples were denatured in Laemmlli buffer (63 mM Tris pH 6.8, 2% SDS, 10% glycerol, 1 mg/ml bromophenol blue, 1% β-mercaptoethanol) at 95°C for five minutes. To separate proteins based on molecular weight, the proteins were subjected to SDS polyacrylamide gel electrophoresis and transferred to nitrocelluose membrane by semi-dry transfer. Nonspecific antibody binding was blocked by incubation with Phosphate buffer saline with 0.05% Tween20 (PBS-T), containing 5% dry milk for 30 minutes at RT. Following the incubation of the membranes in primary antibodies (mouse-anti-FLAG or mouse-anti-Pgk, 1:1000 in PBS-T +5% dry milk) at 4°C overnight, the membranes were washed five times in PBS-T and incubated in secondary antibody (goat-anti-mouse HRP-conjugated, 1:4000 in PBS-T+ 5% dry milk) for 45 minutes at RT. Membranes were washed five time in PBS-T, enhanced chemiluminescence solution (Thermo Fischer) was applied and antibody signal was detected with a BioRad Chemidoc MP system. All images were exported as JPEGs using ImageLab6 (BioRad) and cropped in Adobe Photoshop CC. The western blot shows one representative blot from N=3 replicates performed in parallel with the associated metabolomics and MDC assays.

### Extraction of whole cell metabolites from yeast

For analysis of whole cell metabolite analysis, yeast cells were grown for 15-17 hours to a density of 0.3-0.5 x 10^7^ cells/ml and were treated with indicated drugs for 2 hours. For metabolite analysis on untreated cells, yeast were grown 15-17 hours to a density of 0.5-0.9 x 10^7^ cells/ml. 4 x 10^7^ cells/ml were harvested by centrifugation for 3 minutes at 5000g, washed once with water and cell pellets were shock frozen in liquid nitrogen.

Whole cell metabolites were extracted from yeast cell pellets as previously described, with slight modifications (Canelas et al., 2009). Briefly, 0.4 μg of the internal standard succinic-d4 acid was added to each sample. Next, 1ml of boiling 75% EtOH was added to each pellet, followed by vortex mixing and incubation at 90°C for 3 minutes with intermittent vortex mixing. Cell debris were removed by centrifugation at 7000g for 5 minutes at -10°C. Supernatants were transferred to new tubes and dried *en vacuo.* Process blank samples were made using only extraction solvent and no cell culture pellet.

### GC-MS analysis

GC-MS analysis was performed with an Agilent 5977b GC-MS MSD-HES fit with an Agilent 7693A automatic liquid sampler. Dried samples were suspended in 40 µL of a 40 mg/mL O-methoxylamine hydrochloride (MOX) (MP Bio #155405) in dry pyridine (EMD Millipore #PX2012-7) and incubated for one hour at 37 °C in a sand bath. 25 µL of this solution was added to auto sampler vials.10 µL from the remaining solution for every sample was used to create pooled QC and 25 µL of pooled QC was added to auto sampler vials. 60 µL of N-methyl-N-trimethylsilyltrifluoracetamide (MSTFA with 1% TMCS, Thermo #TS48913) was added automatically via the auto sampler and incubated for 30 minutes at 37 °C. After incubation, samples were vortexed and 1 µL of the prepared sample was injected into the gas chromatograph inlet in the split mode with the inlet temperature held at 250 °C. A 5:1 split ratio was used for analysis for the majority of metabolites. Any metabolites that saturated the instrument at the 5:1 split were analyzed at a 50:1 split ratio. The gas chromatograph had an initial temperature of 60 °C for one minute followed by a 10 °C/min ramp to 325 °C and a hold time of 10 minutes. A 30-meter Agilent Zorbax DB-5MS with 10 m Duraguard capillary column was employed for chromatographic separation. Helium was used as the carrier gas at a rate of 1 mL/min.

Data was collected using MassHunter software (Agilent). Metabolites were identified and their peak area was recorded using MassHunter Quant. This data was transferred to an Excel spread sheet (Microsoft, Redmond WA). Metabolite identity was established using a combination of an in-house metabolite library developed using pure purchased standards, the NIST library and the Fiehn library. Values for each metabolite were normalized to the internal standard in each sample, normalized to sum and are displayed as fold change compared to the control sample. Data was analyzed using the in-house ‘MetaboAnalyst’ software tool. All steady-state metabolite data is provided in Table S5.

## QUANTIFICATION AND STATISTICAL ANALYSIS

All experiments were repeated at least 3 times and all attempts at replication were successful. For yeast MDC assays, N=3 replicates for n=100 cells were used for quantification and statistical analysis. For whole cell metabolite and lipid analysis, N=three to four biological replicates were analyzed in the same run. All statistical analysis was performed in Prism (Graphpad) and the used statistical test is indicated in the corresponding figure legend. No data were excluded from the analysis with the exception of some metabolomics samples that did not meet the quality control cutoff. In the latter case, all samples of the affected biological replicate were excluded from any further analysis. No randomization or blinding was used as all experiments were performed with defined laboratory reagents and yeast strains of known genotypes.

## ONLINE SUPPLEMENTAL MATERIAL

Fig. S1 shows representative widefield images and corresponding line scan analysis of internalization of Bap2 tagged GFP upon different growth and treatment conditions as indicated. Fig. S2 provides additional analysis showing that retrograde pathway mutants control MDC formation. Table S1 lists the yeast strains used in this study. Table S2 lists the oligonucleotides used in this study. Table S3 lists bacterial strains, chemicals, antibodies, plasmids, and software used in this study. Table S4 contains all RNA sequencing data. Table S5 contains all metabolomics data.

## DATA AVAILABILITY

All reagents used in this study are available upon request. All other data reported in this paper will be shared by the lead contact upon request. This paper does not report original code. Any additional information required to reanalyze the data reported in this paper is available from the lead contact upon request.

## ACKNOWLEDGEMENTS

We thank past and present members of the A.L. Hughes group for discussion and manuscript comments. We thank members of Janet. M. Shaw laboratory for providing reagents, mitochondrial antibodies, and support early on in the project. Research was supported by National Institutes of Health grants GM119694 and AG061376 to A.L.H. Metabolomics analysis was performed at the Metabolomics Core Facility at the University of Utah. Mass spectrometry equipment was obtained through NCRR Shared Instrumentation Grant 1S10OD016232-01, 1S10OD018210-01A1 and 1S10OD021505-01. We acknowledge the Metabolomics Core Facility at the University of Utah for use of Gas chromatography Mass Spectrometry (GC-MS) equipment and thank Quentinn Pearce and James Cox for their assistance in GC-MS instrument running and metabolomics analysis. Additionally, for RNA sequencing and corresponding analysis, research reported in this publication utilised the High-Throughput Genomics and Cancer Bioinformatics Shared Resource at Huntsman Cancer Institute at the University of Utah and was supported by the National Cancer Institute of the National Institutes of Health under Award Number P30CA042014. The content is solely the responsibility of the authors and does not necessarily represent the official views of the NIH.

## AUTHOR CONTRIBUTIONS

Conceptualization, N.R. and A.L.H.; methodology, N.R.; formal analysis, N.R.; investigation, N.R.; writing – original draft, N.R.; writing – review and editing, N.R. and A.L.H.; visualization, N.R.; supervision, A.L.H.; funding acquisition, A.L.H.

## CONTACT FOR REAGENT AND RESOURCE SHARING

Further information and requests for resources and reagents should be directed to and will be fulfilled by the Lead Contact, Adam Hughes. All unique/stable reagents generated in this study are available from the Lead Contact without restrictions.

---

**Figure S1.**
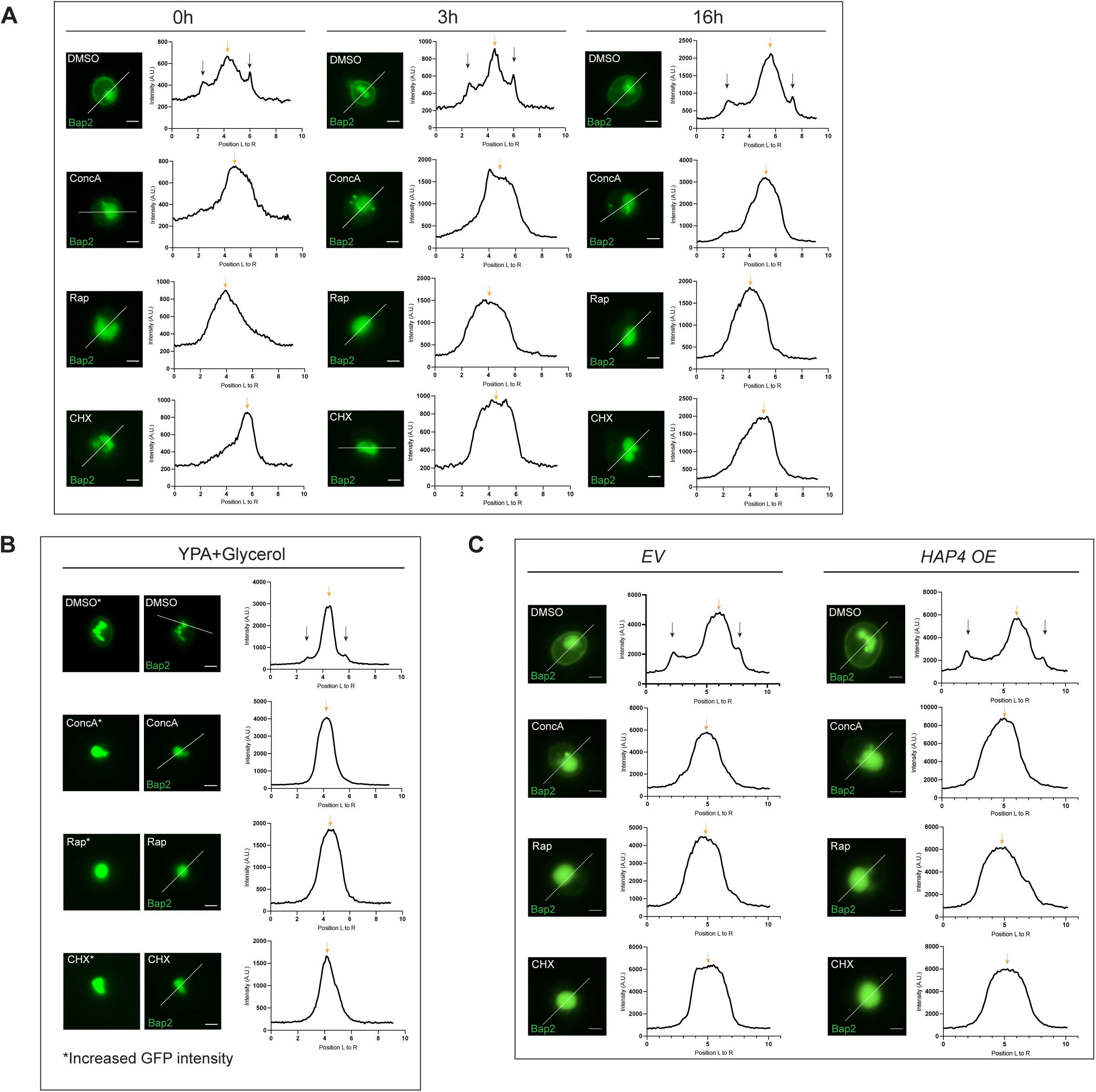
Respiratory state does not impact plasma membrane nutrient transporter remodeling (Related to. **figures 1 and 2).** (A) Widefield microscopy images and corresponding line scan analysis showing internalization of Bap2-GFP upon treatment with concA, rap or cHX. Yellow arrow indicates vacuolar GFP, black arrow indicates GFP on plasma membrane. Scale bar, 2μm. (B) Widefield microscopy images and corresponding line scan analysis showing internalization of Bap2-GFP upon treatment with concA, rap or cHX. Yellow arrow indicates vacuolar GFP, black arrow indicates GFP on plasma membrane. Left image (*) shows where fluorescence intensity has been increased post-imaging to visualize the internalization of Bap2-GFP upon treatment. Scale bar, 2μm. (C) Widefield microscopy images and corresponding line scan analysis showing internalization of Bap2-GFP in *HAP4* overexpressing or *empty vector (EV)* control cells upon treatment with concA, rap or cHX. Yellow arrow indicates vacuolar GFP, black arrow indicates GFP on plasma membrane.

**Figure S2.**
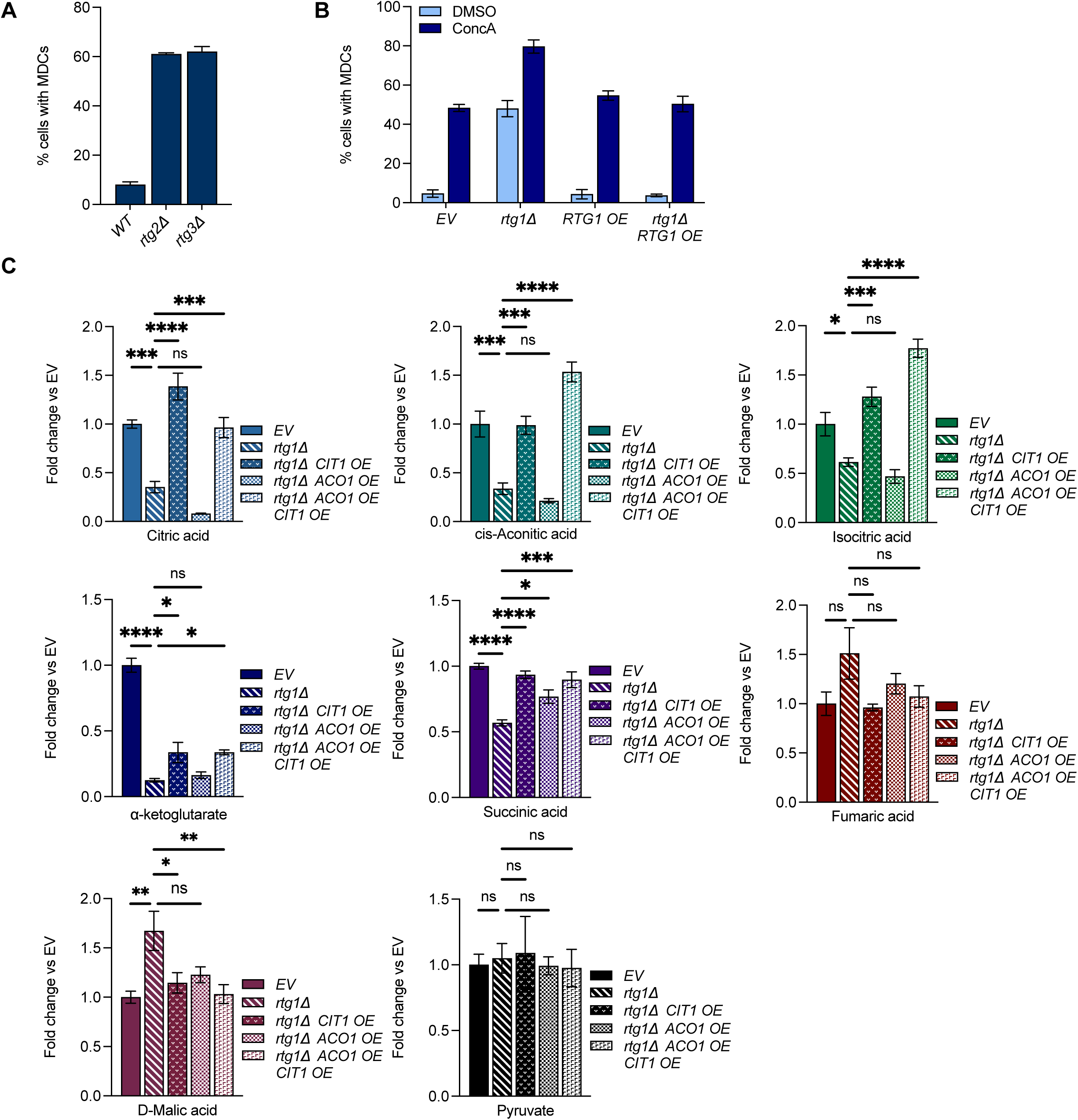
Retrograde pathway mutants control MDC formation (Related to. **figure 5).** (A) Quantification of M+ DC formation in *WT* and indicated mutant cells. Error bars show +/- SEM for N=3 replicates with n=100 cells per replicate. (B) Quantification of MDC formation in *EV* and indicated mutant cells, treated with DMSO or concA. Error bars show +/- SEM for N=3 replicates with n=100 cells per replicate. (C) Analysis of whole cell pyruvate and TCA cycle metabolite levels in *EV*, *rtg1*1Δ** and *rtg1*1Δ** overexpressing *CIT1, ACO1* or *CIT1* and *ACO1* mutant cells grown in high amino acid medium. Error bars show +/- SEM for N=3 replicates. Statistical comparison shows difference to the *rtg1*1Δ** mutant cells. n.s., not significant, *p<0.0332, **p<0.0021, ***p<0.0002, ****<0.0001, Two-way ANOVA with Šídák’s multiple comparisons test.

## REFERENCES

1. Alan, L., Scorrano, L., 2022. Shaping fuel utilization by mitochondria. Curr. Biol. 32, R618– R623. 10.1016/j.cub.2022.05.006

2. Baker Brachmann, C., Davies, A., Cost, G.J., Caputo, E., Li, J., Hieter, P., Boeke, J.D., 1998. Designer deletion strains derived fromSaccharomyces cerevisiae S288C: A useful set of strains and plasmids for PCR-mediated gene disruption and other applications. Yeast 14, 115–132. 10.1002/(SICI)1097-0061(19980130)14:2&lt;115::AID-YEA204ti3.0.CO;2-2

3. Butow, R.A., Avadhani, N.G., 2004. Mitochondrial Signaling:The Retrograde Response. Mol. Cell 14, 1–15. 10.1016/S1097-2765(04)00179-0

4. Canelas, A.B., Ten Pierick, A., Ras, C., Seifar, R.M., Van Dam, J.C., Van Gulik, W.M., Heijnen, J.J., 2009. Quantitative Evaluation of Intracellular Metabolite Extraction Techniques for Yeast Metabolomics. Anal. Chem. 81, 7379–7389. 10.1021/ac900999t

5. Chandel, N.S., 2014. Mitochondria as signaling organelles. BMC Biol. 12, 34. 10.1186/1741-7007-12-34

6. Chen, E.Y., Tan, C.M., Kou, Y., Duan, Q., Wang, Z., Meirelles, G.V., Clark, N.R., Ma’ayan, A., 2013. Enrichr: interactive and collaborative HTML5 gene list enrichment analysis tool. BMC Bioinformatics 14, 128. 10.1186/1471-2105-14-128

7. De Deken, R.H., 1966. The Crabtree Effect: A Regulatory System in Yeast. J. Gen. Microbiol. 44, 149–156. 10.1099/00221287-44-2-149

8. Dobin, A., Davis, C.A., Schlesinger, F., Drenkow, J., Zaleski, C., Jha, S., Batut, P., Chaisson, M., Gingeras, T.R., 2013. STAR: ultrafast universal RNA-seq aligner. Bioinformatics 29, 15–21. 10.1093/bioinformatics/bts635

9. Egner, A., Jakobs, S., Hell, S.W., 2002. Fast 100-nm resolution three-dimensional microscope reveals structural plasticity of mitochondria in live yeast. Proc. Natl. Acad. Sci. 99, 3370–3375. 10.1073/pnas.052545099

10. English, A.M., Kornmann, B., Shaw, J.M., Hughes, A.L., 2020. ER-Mitochondria Contacts Promote Mitochondrial-Derived Compartment Biogenesis (preprint). Cell Biology. 10.1101/2020.03.13.991133

11. Epstein, C.B., Waddle, J.A., Hale, W., Davé, V., Thornton, J., Macatee, T.L., Garner, H.R., Butow, R.A., 2001. Genome-wide Responses to Mitochondrial Dysfunction. Mol. Biol. Cell 12, 297–308. 10.1091/mbc.12.2.297

12. Ewels, P., Magnusson, M., Lundin, S., Käller, M., 2016. MultiQC: summarize analysis results for multiple tools and samples in a single report. Bioinformatics 32, 3047–3048. 10.1093/bioinformatics/btw354

13. Grauslund, M., Didion, T., Kielland-Brandt, M.C., Andersen, H.A., 1995. BAP2, a gene encoding a permease for branched-chain amino acids in Saccharomyces cerevisiae. Biochim. Biophys. Acta BBA - Mol. Cell Res. 1269, 275–280. 10.1016/0167-4889(95)00138-8

14. Hatakeyama, R., De Virgilio, C., 2019. TORC1 specifically inhibits microautophagy through ESCRT-0. Curr. Genet. 65, 1243–1249. 10.1007/s00294-019-00982-y

15. Hazelwood, L.A., Daran, J.-M., van Maris, A.J.A., Pronk, J.T., Dickinson, J.R., 2008. The Ehrlich Pathway for Fusel Alcohol Production: a Century of Research on Saccharomyces cerevisiae Metabolism. Appl. Environ. Microbiol. 74, 2259–2266. 10.1128/AEM.02625-07

16. Hughes, A.L., GoIschling, D.E., 2012. An early age increase in vacuolar pH limits mitochondrial function and lifespan in yeast. Nature 492, 261–265. 10.1038/nature11654

17. Hughes, A.L., Hughes, C.E., Henderson, K.A., Yazvenko, N., GoIschling, D.E., 2016. Selective sorting and destruction of mitochondrial membrane proteins in aged yeast. eLife 5, e13943. 10.7554/eLife.13943

18. Jazwinski, S.M., Kriete, A., 2012. The Yeast Retrograde Response as a Model of Intracellular Signaling of Mitochondrial Dysfunction. Front. Physiol. 3. 10.3389/fphys.2012.00139

19. Karbowski, M., Youle, R.J., 2011. Regulating mitochondrial outer membrane proteins by ubiquitination and proteasomal degradation. Curr. Opin. Cell Biol. 23, 476–482. 10.1016/j.ceb.2011.05.007

20. Klein, C.J.L., Olsson, L., Nielsen, J., 1998. Glucose control in Saccharomyces cerevisiae: the role of MIG1 in metabolic functions. Microbiology 144, 13–24. 10.1099/00221287-144-1-13

21. König, T., McBride, H.M., 2024. Mitochondrial-derived vesicles in metabolism, disease, and aging. Cell Metab. 36, 21–35. 10.1016/j.cmet.2023.11.014

22. Labbé, K., Murley, A., Nunnari, J., 2014. Determinants and Functions of Mitochondrial Behavior. Annu. Rev. Cell Dev. Biol. 30, 357–391. 10.1146/annurev-cellbio-101011-155756

23. Lascaris, R., Bussemaker, H.J., Boorsma, A., Piper, M., van der Spek, H., Grivell, L., Blom, J., 2002. [No title found]. Genome Biol. 4, R3. 10.1186/gb-2002-4-1-r3

24. Li, X., Straub, J., Medeiros, T.C., Mehra, C., Den Brave, F., Peker, E., Atanassov, I., Stillger, K., Michaelis, J.B., Burbridge, E., Adrain, C., Münch, C., Riemer, J., Becker, T., Pernas, L.F., 2022. Mitochondria shed their outer membrane in response to infection-induced stress. Science 375, eabi4343. 10.1126/science.abi4343

25. Liao, Y., Smyth, G.K., Shi, W., 2014. featureCounts: an efficient general purpose program for assigning sequence reads to genomic features. Bioinformatics 30, 923–930. 10.1093/bioinformatics/bI656

26. Lin, C.H., MacGurn, J.A., Chu, T., Stefan, C.J., Emr, S.D., 2008. Arrestin-Related Ubiquitin- Ligase Adaptors Regulate Endocytosis and Protein Turnover at the Cell Surface. Cell 135, 714–725. 10.1016/j.cell.2008.09.025

27. Liu, Z., Butow, R.A., 2006. Mitochondrial Retrograde Signaling. Annu. Rev. Genet. 40, 159–185. 10.1146/annurev.genet.40.110405.090613

28. Liu, Z., Butow, R.A., 1999. A Transcriptional Switch in the Expression of Yeast Tricarboxylic Acid Cycle Genes in Response to a Reduction or Loss of Respiratory Function. Mol. Cell. Biol. 19, 6720–6728. 10.1128/MCB.19.10.6720

29. Love, M.I., Huber, W., Anders, S., 2014. Moderated estimation of fold change and dispersion for RNA-seq data with DESeq2. Genome Biol. 15, 550. 10.1186/s13059-014-0550-8

30. Martin, M., 2011. Cutadapt removes adapter sequences from high-throughput sequencing reads. EMBnet.journal 17, 10. 10.14806/ej.17.1.200

31. Morawska, M., Ulrich, H.D., 2013. An expanded tool kit for the auxin-inducible degron system in budding yeast. Yeast Chichester Engl. 30, 341–351. 10.1002/yea.2967

32. Mülleder, M., Capuano, F., Pir, P., Christen, S., Sauer, U., Oliver, S.G., Ralser, M., 2012. A prototrophic deletion mutant collection for yeast metabolomics and systems biology. Nat. Biotechnol. 30, 1176–1178. 10.1038/nbt.2442

33. Nikko, E., Pelham, H.R.B., 2009. Arrestin-Mediated Endocytosis of Yeast Plasma Membrane Transporters. Traffic 10, 1856–1867. 10.1111/j.1600-0854.2009.00990.x

34. Nunnari, J., Suomalainen, A., 2012. Mitochondria: In Sickness and in Health. Cell 148, 1145– 1159. 10.1016/j.cell.2012.02.035

35. Parnell, E.J., Jenson, E., Miller, M.P., 2023. An interaction hub on Ndc80 complex facilitates dynamic recruitment of Mps1 to yeast kinetochores to promote accurate chromosome segregation (preprint). Cell Biology. 10.1101/2023.11.07.566082

36. Pickles, S., Vigié, P., Youle, R.J., 2018. Mitophagy and Quality Control Mechanisms in Mitochondrial Maintenance. Curr. Biol. 28, R170–R185. 10.1016/j.cub.2018.01.004

37. Quirós, P.M., Langer, T., López-Oxn, C., 2015. New roles for mitochondrial proteases in health, ageing and disease. Nat. Rev. Mol. Cell Biol. 16, 345–359. 10.1038/nrm3984

38. Ravanelli, S., Den Brave, F., Hoppe, T., 2020. Mitochondrial Quality Control Governed by Ubiquitin. Front. Cell Dev. Biol. 8, 270. 10.3389/fcell.2020.00270

39. RuIer, J., Hughes, A.L., 2015. Power2: The power of yeast genetics applied to the powerhouse of the cell. Trends Endocrinol. Metab. 26, 59–68. 10.1016/j.tem.2014.12.002

40. Schindelin, J., Arganda-Carreras, I., Frise, E., Kaynig, V., Longair, M., Pietzsch, T., Preibisch, S., Rueden, C., Saalfeld, S., Schmid, B., Tinevez, J.-Y., White, D.J., Hartenstein, V., Eliceiri, K., Tomancak, P., Cardona, A., 2012. Fiji: an open-source playorm for biological-image analysis. Nat. Methods 9, 676–682. 10.1038/nmeth.2019

41. Schuler, M.-H., English, A.M., Xiao, T., Campbell, T.J., Shaw, J.M., Hughes, A.L., 2021. Mitochondrial-derived compartments facilitate cellular adaptation to amino acid stress. Mol. Cell 81, 3786–3802.e13. 10.1016/j.molcel.2021.08.021

42. Sheff, M.A., Thorn, K.S., 2004. Optimized casseIes for Fluorescent protein tagging in Saccharomyces cerevisiae. Yeast 21, 661–670. 10.1002/yea.113010.1002/yea.1130

43. Shpilka, T., Haynes, C.M., 2018. The mitochondrial UPR: mechanisms, physiological functions and implications in ageing. Nat. Rev. Mol. Cell Biol. 19, 109–120. 10.1038/nrm.2017.110

44. Sikorski, R.S., Hieter, P., 1989. A system of shuIle vectors and yeast host strains designed for efficient manipulation of DNA in Saccharomyces cerevisiae. Genetics 122, 19–27. 10.1093/genetics/122.1.19

45. Spinelli, J.B., Haigis, M.C., 2018. The multifaceted contributions of mitochondria to cellular metabolism. Nat. Cell Biol. 20, 745–754. 10.1038/s41556-018-0124-1

46. Sugiura, A., McLelland, G., Fon, E.A., McBride, H.M., 2014. A new pathway for mitochondrial quality control: mitochondrial-derived vesicles. EMBO J. 33, 2142–2156. 10.15252/embj.201488104

47. Wallace, D.C., 2005. A Mitochondrial Paradigm of Metabolic and Degenerative Diseases, Aging, and Cancer: A Dawn for Evolutionary Medicine. Annu. Rev. Genet. 39, 359–407. 10.1146/annurev.genet.39.110304.095751

48. Westermann, B., 2012. Bioenergetic role of mitochondrial fusion and fission. Biochim. Biophys. Acta BBA - Bioenerg. 1817, 1833–1838. 10.1016/j.bbabio.2012.02.033

49. Wilson, Z.N., Balasubramaniam, S.S., Wopat, M., Hughes, A.L., 2023a. Mitochondrial-Derived Compartments Remove Surplus Proteins from the Outer Mitochondrial Membrane (preprint). Cell Biology. 10.1101/2023.07.07.548175

50. Wilson, Z.N., West, M., English, A.M., Odorizzi, G., Hughes, A.L., 2023b. Mitochondrial- Derived Compartments are Multilamellar Domains that Encase Membrane Cargo and Cytosol (preprint). Cell Biology. 10.1101/2023.07.07.548169

51. Xiao, T., English, A.M., Wilson, Z.N., Maschek, J.Alan., Cox, J.E., Hughes, A.L., 2024. The phospholipids cardiolipin and phosphatidylethanolamine differentially regulate MDC biogenesis. J. Cell Biol. 223, e202302069. 10.1083/jcb.202302069

52. Xu, Z., Wei, W., Gagneur, J., Perocchi, F., Clauder-Münster, S., Camblong, J., Guffanti, E., Stutz, F., Huber, W., Steinmetz, L.M., 2009. Bidirectional promoters generate pervasive transcription in yeast. Nature 457, 1033–1037. 10.1038/nature07728

53. Youle, R.J., Van Der Bliek, A.M., 2012. Mitochondrial Fission, Fusion, and Stress. Science 337, 1062–1065. 10.1126/science.1219855

54. Zecchini, V., Paupe, V., Herranz-Montoya, I., Janssen, J., Wortel, I.M.N., Morris, J.L., Ferguson, A., Chowdury, S.R., Segarra-Mondejar, M., Costa, A.S.H., Pereira, G.C., Tronci, L., Young, T., Nikitopoulou, E., Yang, M., Bihary, D., Caicci, F., Nagashima, S., Speed, A., Bokea, K., Baig, Z., Samarajiwa, S., Tran, M., Mitchell, T., Johnson, M., Prudent, J., Frezza, C., 2023. Fumarate induces vesicular release of mtDNA to drive innate immunity. Nature 615, 499–506. 10.1038/s41586-023-05770-w

